# Predicting Suitable Habitats Of Critically Endangered Chinese Pangolin In Assam, India

**DOI:** 10.1101/2023.10.05.561022

**Authors:** Kuladip Sarma, Murali Krishna, Barin Kumar Boro, Malabika Kakati Saikia, Bidyut Sarania, Himolin Basumatary, Bhrigu Prasad Saikia, Prasanta Kumar Saikia

## Abstract

Chinese pangolin is a least known species in terms of its ecology and population status in India. The present study of Chinese pangolin in Kaziranga-Karbi-Anglong landscape aimed to gather information on species distribution and population status and also an account of threats of the species was investigated. A total of 24 occurrence points were collected from all over Assam and used to get a predictive potential distribution map for the species via Maxent and Random Forest models. To understand the knowledge level of the local people, a total of 13 villages in the north east boundary of North Karbi Anglong Wildlife Sactuary have been surveyed and 160 respondents were interviewed. The AUC values for the two models namely Maxent (0.736) and Random Forest (0.87) were different. Out of the six environmental parameters chosen, the altitude clarified the maximum variation in the model (62 %) followed by the seasonal temperature (bio4; 19.6 %) in the Maxent model. At the same time, these two variables were also found to be of greater significance in the Random Forest model. The species has been least seen by most of the respondents in the last 10 years (43.7%). Most of the respondent of age class 18-24 years never saw a pangolin (82.5%) in the area which depict the population decline of the spcies. Scales were the most used body parts for traditional medicine (58.8%) especially for exceesive saliva secretion in children. The resultant potential habitat in the predictive distribution map in Assam needs to be confirmed with ground verification and accordingly inclusion in the priority conservation zone for Chinese pangolin in Assam should be done.

## Introduction

Chinese pangolin (*Manis pentadactyla*) one of the four species that occurs in Indo-Malayan biogeographic region distributed in eastern Himalayan parts of Nepal, Bhutan, northeast, India and Southeast China (Yang et al., 2018; Sharma et al., 2020; Dorji et al., 2020; Kumar et al., 2020). The distribution of the *M. pentadactyla* overlaps with the Indian pangolin (*Manis crassicaudata*), whereas sunda pangolin (*Manis javanica*) and Philippine pangolin (*Manis culionensis*) have restricted distribution in Southeast Asia and Philippine respectively (Challender, 2020). The pangolin species from Indo-Malayan region belongs to the genera of Manis, and three out of the four species of the region are categorized as critically endangered including chinese pangolin (IUCN, 2020). The population of these species are drastically declining due to the extensive hunting for meat, habitat loss and used of different body parts in traditional medicine across its distribution range. Moreover, *M. pentadactyla* is reported as a one of the most trafficking mammals of the world (Challender et al., 2020). Kumar et al. (2020) reported that northeast India is the gateway of illegal trafficking of the species to the China and Myanmar.

The knowledge on distribution ecology of *M. pentadactyla* is limited except few studies in Nepal, Bhutan and China have been reported till date (Bhandari and Chalise, 2014; Thapa et al., 2018; Sharma et al., 2020; Nash et al., 2016; Yang et al., 2018). However, there is no study on distribution ecology of the species from the north-eastern region of India (D’cruz et al., 2018). The altitudinal distribution ranges of the species are reported from 500 m above mean seal level (amsl) to 2406m amsl in eastern Himalaya biogeographic region (Dorji et al., 2020). Suwal et al. (2020) study reported that ground and canopy cover are importance aspects for the occurrence of the species. Northeast India is a potential population threshold of *M. pentadactyla*; at the same time, this region is also most vulnerable for the species due to extensive hunting and rapid loss of forest cover (Kumar et al., 2020). The state of Assam of northeast India (8 states) is the tropical habitat region of *M. pentadactyla* (Sarma, 2015). However, the knowledge on habitat distribution pattern as well as the status of the habitat is significantly limited from the state. There is a difficulty in detection of pangolin using the generalize methodology that are best fit for ground dwelling mammals other than pangolin. Most of studies on the assessment of population rely on the secondary data including the associated fauna and habitat condition, information on community perception (Nash et al., 2016), market and trade, and trafficking dynamics (Newton et al., 2008).

One of the options for the study of potential habitat of the species is predictive species distribution modeling (SDM). SDM is a very useful approach for determining, in a reliable way, areas where suitable environmental conditions exist for species and where future exploration would be fruitful (Phillips et al., 2004; Dudík et al,. 2007). MaxEnt (available for download from http://www.cs.princeton.edu/*schapire/maxent/) modeling, which is based on the principle of maximum entropy is mostly used SDM for chinese pangolin which is useful tool for determining the potential distribution of species (Ortega-Huerta and Townsend, 2008). Maximum entropy (Maxent) model (Phillips et al., 2004) is based on statistical mechanics (Jaynes, 1957) and its general purpose is predicting the potential distribution of species. The method requires only species occurrence data and environmental information (Elith *et al*., 2011). Another major advantage of the model includes allowing the use of both continuous and categorical data, which incorporates the interactions between the predictor variables as well (Phillips et al., 2006). Maxent distribution model has been widely used in the prediction of non-human primate distribution, plant distribution and distributions of other wild animals (Sarania et al., 2017; Wan, 2019). More recently, Maxent is used for *M. pentadactyla* to predict the suitable habitat in Nepal and Bhutan (Suwal et al., 2020; Thapa et al., 2018). Moreover, other machine learning techniques such as Random Forest Classification has also been proved to be competent in predicting suitable habitat of wildlife with limited occurrence information. Although, it is obscure to find out the best model for prediction of suitable habitat, Maxent and Random forest have given best result in terms of model evaluation (Duarte et al., 2018). Therefore, it is recommended to average the output of more than one model to get the final predictive map, rather considering a single model for comparison.

The limited understanding on the ecological niche of critically endanger *M. pentadactyla* in Assam is hindering in population survey and obstructed in identifying areas; where urgent intervene is required to safeguard the species. Thus, we carried out a study on potential habitat distribution of *M. pentadactyla* and assessed the habitat status (tree canopy cover as proxy for habitat condition). We used Maxent and Random Forest model, averaged the output to map the potential habitat range of *M. pentadactyla*, and assessed the tree cover change (two decades) using Google earth engine.

## Methods

### Study area

The distribution survey has been conducted to the south of Kaziranga National Park in Karbi Anglong districts, Assam. The landscape is very unique being in the junction of hill area in Karbi Anglong and flood plains in Kaziranga National Parkand thus, rich in faunal and floral components. The Karbi Anglong area is well protected with establishment of numbers of WLS and Reserve Forests etc. The landscape is the mosaic of human-forest interactions. The selection of the site is based on the information of chinese pangolin captured and killed/release in the past 10 years (only published news articles in local daily news papers). As all the three sites are protected under WPA, 1972, and having well managed forest staff, our target will be to encompass local people in the greater umbrella of the project to gather both local ecological information as well as to aware them about the importance of the species. From the conservation point of view, the present study is less focused especially in terms of species conservation perspective. Moreover, the ethnic tribes live in and around the study area are having most diverse cultural and traditional knowledge system.

### Distribution survey

Survey was conducted in Karbi Anglong hills and adjoining areas towards Kaziranga National Park, Assam from 2015 to 2017 to know the distribution pattern of chinese pangolin *(Manis pentadactyla)*. As the species distribution range is least studied, various information regarding the species’ occurrence were collected from different sources as well as from the direct sightings of the burrows of the species across the state of Assam. Local knowledge has been given top priority during the survey to collect information about the habitat of the species and other information such as hunting, use of animal parts in traditional medicine etc.GPS locations of known burrows of the species were used as preliminary occurrence data to run the distribution model along with other relevant environmental data.

### Environmental layers and occurrence data

Twenty environmental layers including temperature, precipitation, etc. and elevation information (SRTM) were downloaded from the WorldClim website (www.worldclim.org) for 30 seconds of spatial resolution. Although bioclimatic variables are important to explain the distribution of species (Busby 1986, Nix 1986), for modeling if the explanatory variables are used without screening it may inadvertently lead to the inclusion of those variables which are also highly correlated and basically have the same set of information. In such cases, these variables may be inclining statistical weight toward themselves, and when this happens, it may undesirably result in over-fitting of the model (Anderson and Gonzales, 2011; van Gils et al., 2014). Data of occurrence of *M. pentadactyla* was gathered through extensive field surveys conducted across Assam, India. A total of 24 independent localities were collected from the field, and all localities were used in the final modeling process.

The variance inflation factor (VIF) from the package usdm under R v 4.0.2 (R Development Core Team, 2017) was used to screen the variables for collinearity and variables having VIF>10 have been removed from the model as strong collinearity affects model performance (Quinn & Keough, 2002). There are various reports available where VIF has been applied in several species distribution models (Lauria et al., 2015; Ranjitkar et al., 2014a; Ranjitkar et al., 2014b).Thus, we have selected 5 statistical and biologically meaningful bioclimatic variables (Table 2) along with the altitude layer for modeling habitat distribution for the *M. pentadactyla*.

**Table 1.** Distribution data (occurrence data) of Chinese Pangolin in Assam used in the predictive distribution model of the species.

| Sl./No. | Name of Locations | Longitude (E) | Latitude (N) | Source |
| --- | --- | --- | --- | --- |
| 1 | Rani Picnic spot | 91.62886 | 26.03469 | Primary |
| 2 | Rani Picnic spot | 91.61597 | 26.02711 | Primary |
| 3 | Manas National Park | 91.21319 | 26.76703 | Primary |
| 4 | Manas National Park | 91.23268 | 26.76824 | Primary |
| 5 | Ranga De-reserve | 93.83516 | 27.08477 | Secondary |
| 6 | Hoollongapar Gibbon WLS | 94.35483 | 26.6691 | Secondary |
| 7 | Hoollongapar Gibbon WLS | 94.33628 | 26.68485 | Secondary |
| 8 | Behali Reserve Forest | 93.28066 | 26.86707 | Secondary |
| 9 | Kuthori | 93.24772 | 26.56089 | Primary<br>(Kaziranga-KarbiAnglong<br>Landscape) |
| 10 | Dolamari | 93.56245 | 26.5379 |  |
| 11 | Diphu | 93.33578 | 25.87144 |  |
| 12 | Lumding | 93.19962 | 25.7911 |  |
| 13 | Bokolia | 93.29798 | 25.97595 |  |
| 14 | Panbari Reserve Forest | 93.54457 | 26.62173 |  |
| 15 | Nellie | 92.3179 | 26.07845 | Secondary |
| 16 | Pobitora WLS | 92.05225 | 26.2225 | Secondary |
| 17 | Akashi Ganga | 92.93582 | 26.18016 | Secondary |
| 18 | Pavo RF | 95.27994 | 27.83781 | Secondary |
| 19 | Dulung RF | 94.17902 | 27.4246 | Secondary |
| 20 | Ranga RF | 93.99745 | 27.27478 | Secondary |
| 21 | Namti | 94.65112 | 26.74971 | Secondary |
| 22 | Bogaikhas | 91.2886 | 25.88074 | Primary |
| 23 | Kanamakra | 90.64176 | 26.69049 | Primary |
| 24 | Subonkhata | 91.35973 | 26.77384 | Primary |

Primary=collected by CLP team members
Secondary=collected by volunteers

**Table 2:** Variance Inflation Factor (VIF) of selected bioclimatic variables downloaded from worldclim database.

| Variable Name | Code | Variance Inflation Factor (VIF) |
| --- | --- | --- |
| Isothermality ($(\text{Bio2}/\text{Bio7}) \times 100$) | bio3 | 3.83 |
| Temperature seasonality (standard deviation $\times 100$) | bio4 | 9.90 |
| Mean temperature of wettest quarter | bio8 | 4.40 |
| Precipitation of driest period | bio14 | 7.42 |
| Precipitation seasonality | bio15 | 3.52 |

### Model algorithm

#### Maxent

Maxent (Maximum Entropy) modeling method is one of the most robust and superior bioclimatic modeling approaches for presence-onlydata (Elith et al., 2006, 2011; Wisz, Tamstorf, Madsen, & Jespersen, 2008). The function in Maxent uses environmental data for known-presence locations and for a large number of “background” locations (Hijmans and Elith 2017). The background locations are generated randomly from the raster data from which the environmental data is extracted. The result of this presence only model is a model object which can be used, for example, to predict the suitability of other locations to predict the whole range of a species.

### Random Forest

The system of the Random Forest (Breiman 2001) is an extension of trees of classification and regression (CART; Breiman et al. 1984).In R, random Forest package can be implemented in two different arguments, a predictor variable data.frame and an response vector. If the response variable is a (categorical) factor, random Forest will classify, otherwise it will do regression (Hijmans and Elith2017).

### Model Average Score

In order to determine the best model of the two models, the two model outputs are combined in order to produce a single result, since some authors (e.g. Thuillier 2003) suggested for choosing the best model out of multiple models, model averaging is a simple choice. However, this is a troublesome approach, since the values predicted by the two models are not on the same scale (between 0 and 1), such that we merge the two models with their AUC scores (Hijmans and Elith 2017).

### Model Validation

We selected 75% of the data for training and the rest 25% for testing and 10 percentile threshold rule was employed. AUC (**A**rea **U**nder the receiving operator **C**urve) was used to test the model’s goodness-of-fit. The highest AUC was considered as the best performer. Models have been discriminated based on AUC of receiver operating characteristic (ROC) function (Fielding and Bell 1997). The AUC as threshold-independent measure varies from 0.5 to 1 for complete discrimination of an uninformative model (Elith 2002).AUC is a Rank-Correlation metric. A high AUC suggests, in unbiased data, that sites with high predicted suitability values tend to be areas of known presence and locations with lower model prediction values tend to be areas where the species is not known to be present (absence or a random point). Phillips et al. (2006) outlined the overview of AUC score in the context of presence-only rather than presence/absence data. However, evaluating using AUC has also been criticized (Lobo et al., 2008; Jimfenez-Valverde, 2011), particularly in the context of the spatial extent used to select background points. Generally, higher the degree of extent, the higher is AUC value. Evaluationcriteria for the AUC statistic are as follows: excellent (0.90–1.00), very good (0.8– 0.9), good (0.7–0.8), fair (0.6–0.7), and poor (0.5–0.6) (Swets 1988; Beale and Lennon 2012).

The Kappa coefficient is a further measure of the model prediction results. The Kappa coefficient is based on the maximum value that can better be used in the mixed matrix to calculate the efficiency of the model (Duan et al. 2014).The Kappa statistic assessment criteria are as follows: excellent (0.85-1.0), very good (0.7-0.85), good (0.55-0.7), fair (0.4-0.55), and fail (0.4).The final output was divided into five potential distribution areas which were regrouped with the range of 0 to 1 viz., No Suitability (<0.2); Low Suitability(0.2-0.4); Moderate Suitability (0.4-0.6) and High Suitability (>0.6).

### Gathering local ecological knowledge

Local ecological knowledge was collected through household survey in 100 selected villages in the study area. A target number of ≥10 interviews were conducted per village; this number complies with predicted response saturation levels (White et al., 2005; Guest, 2006). Local village heads and key informants was identified and interviewed through “snowball sampling” (Newing, 2011). Semi-structured questionnaire survey and key informant survey was conducted to assess the existing threats on the species in term of local tratde and medicinal uses during May to August, 2016. A total of thirteen (13) villages in the north east boundary of North Karbi Anglong Wildlife Sactuary have been surveyed and 160 respondents were interviewed.

### Tree canopy cover

We used Hansen Global Forest Cover Change (2000-2019) data set for the assessment of tree canopy cover change in predicted suitable habitat of pangolin. Global Forest Cover (GFC) is a derived product from 30 m spatial resolution based on Landsat data and it has three band viz., treeCover2000, Loss and gain band. The band “tree Cover2000” (Hansen et al., 2013). The band treeCover2000 (consider vegetation taller than 5m height) provides tree canopy cover data for the year of 2000, whereas loss band calculate conversion of forest to non-forest area (2001-2019) and gain band calculate gain of tree canopy cover for the year of 2000-2012. The GFC dataset can be accessed freely on Google Earth Engine platform (GEE) and allows to construct time series for study sites through an Internet-accessible application programming interface of GEE (Gorelick et al., 2017). The GEE has example code for analysis of GFC data set, we modified the code to extract the total annual tree canopy loss area in the predicted habitat for the year of 2001-2019. We used loss band of the product to estimate the loss area of tree canopy cover in the high, moderate and low suitable predicted habitat of *M. pentadactyla*. Subsequently, we analyzed the trend of annual tree canopy loss using Non-parametric Man Mann–Kendall test.

## RESULTS

The AUC values for the two models namely Maxent (0.736) and Random Forest (0.87) were different (Table 3). On the other hand, the kappa coefficient is much higher for Maxent model (0.464) than that of Random Forest (0.013); whereas, the correlation coefficient is higher in Random Forest model (0.186).

**Table 3:** Environmental and its associated variables used in modeling.

| Code | Environmental variables | Unit |
| --- | --- | --- |
| Bio1 | Annual mean temperature | °C |
| Bio2 | Mean diurnal range (mean of monthly max. and min. temp.) | °C |
| Bio3* | Isothermality ((Bio2/Bio7) × 100) |  |
| Bio4 * | Temperature seasonality (standard deviation ×100) |  |
| Bio5 | Maximum temperature of warmest month | °C |
| Bio6 | Minimum temperature of coldest month | °C |
| Bio7 | Temperature annual range (Bio5–Bio6) | °C |
| Bio8* | Mean temperature of wettest quarter | °C |
| Bio9 | Mean temperature of driest quarter | °C |
| Bio10 | Mean temperature of warmest quarter | °C |
| Bio11 | Mean temperature of coldest quarter | °C |
| Bio12 | Annual precipitation | mm |
| Bio13 | Precipitation of wettest period | mm |
| Bio14* | Precipitation of driest period | mm |
| Bio15* | Precipitation seasonality | mm |
| Bio16 | Precipitation of wettest quarter | mm |
| Bio17 | Precipitation of driest quarter | mm |
| Bio18 | Precipitation of warmest quarter | mm |
| Bio19 | Precipitation of coldest quarter | mm |
| Alt* | Elevation | m |
| *Selected variables used in the models. |  |  |

We obtained the variable contribution / importance for both the AUC and Kappa models. Out of the six environmental parameters chosen, the altitude clarified the maximum variation in the model (62 per cent) followed by the seasonal temperature (bio4; 19.6 per cent; Table 4) in the Maxent model. At the same time, these two variables were also found to be of greater significance in the Random Forest model (Gini based importance; score=3.74 and 3.75 respectively).

**Table 4.** Model evaluation statistics for the two models.

| Sl./No. | Model | Statistics |  |  |
| --- | --- | --- | --- | --- |
|  |  | AUC | Kappa | Correlation coefficient (r) |
| 1. | Maxent | 0.736 | 0.464 | 0.068 |
| 2. | Random Forest | 0.87 | 0.013 | 0.186 |

While the form of the absence of presence data used in the Maxent model varies from that seen in Fig.4, there is no statistically meaningful difference (Mann Whitney Rank Test; p>0.05). There were further outliers in the Random Forest model; however, statistical significance was not observed (Mann Whitney Rank Test; p>0.05).

The average of the two models (Figure 2) depicted that the low suitability area is highest (19321.37 km^2^) followed by moderate suitability (2940.95 km^2^) and small area was found under high suitability threshold (24.89 km^2^) (Table 5). While comparing the area statistics of the two models, the area under low suitability is highest in Maxent (23570.96 km^2^) and Random Forest (948.29 km^2^). The average model map shows more area under every type of suitability, primarily in the northern foothills, i.e.Bhutan and Arunachal Pradesh foot hills and south in the foothills of the Mikir hills (Fig. 4).

**Table 5.**
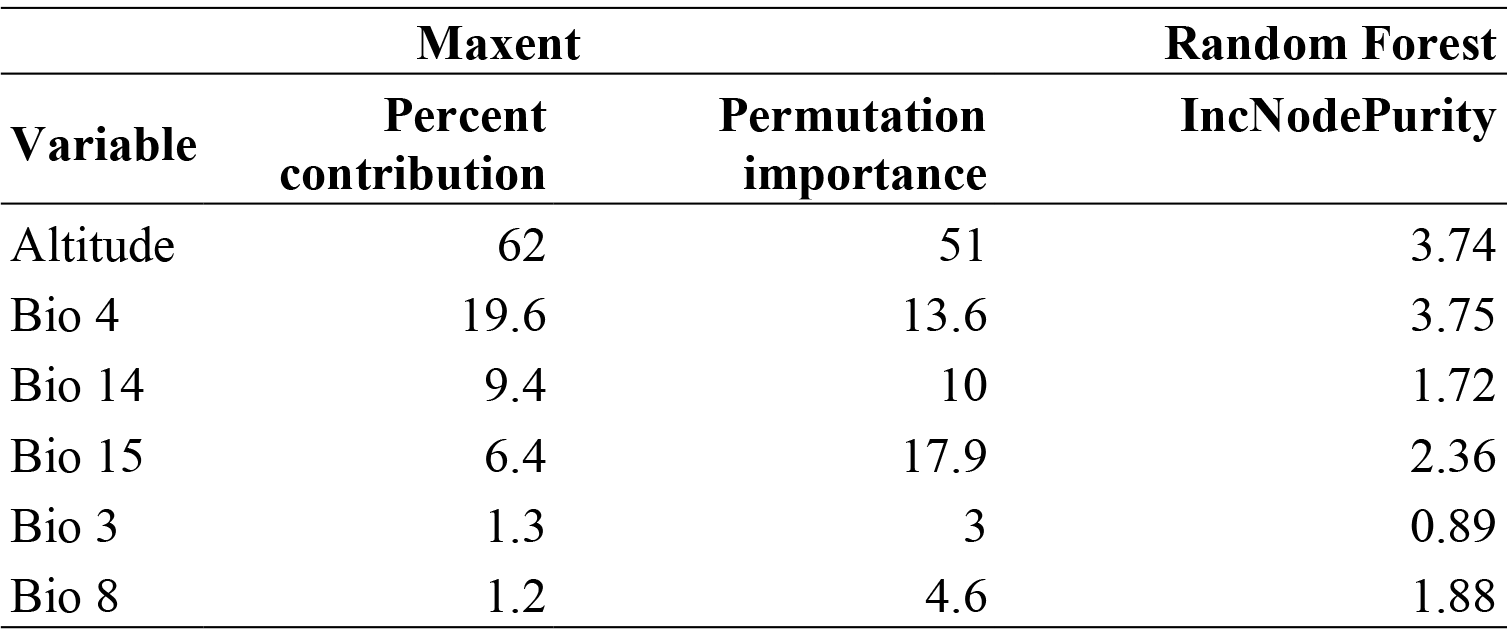
Variable contribution of Maxent and Random Forest model.

| Variable | Maxent |  | Random Forest |
| --- | --- | --- | --- |
|  | Percent contribution | Permutation importance | IncNodePurity |
| Altitude | 62 | 51 | 3.74 |
| Bio 4 | 19.6 | 13.6 | 3.75 |
| Bio 14 | 9.4 | 10 | 1.72 |
| Bio 15 | 6.4 | 17.9 | 2.36 |
| Bio 3 | 1.3 | 3 | 0.89 |
| Bio 8 | 1.2 | 4.6 | 1.88 |

**Table 6.** Area statistics of suitability threshold determined by the models and average score.

| Models | Area (km<sup>2</sup>) |  |  |
| --- | --- | --- | --- |
|  | Low Suitability | Moderate Suitability | High Suitability |
| Maxent | 23570.96 | 11972.25 | 8360.86 |
| Random Forest | 948.29 | 96.29 | 0.76 |
| Model Average | 19321.37 | 2940.95 | 24.89 |

**Fig.1.**
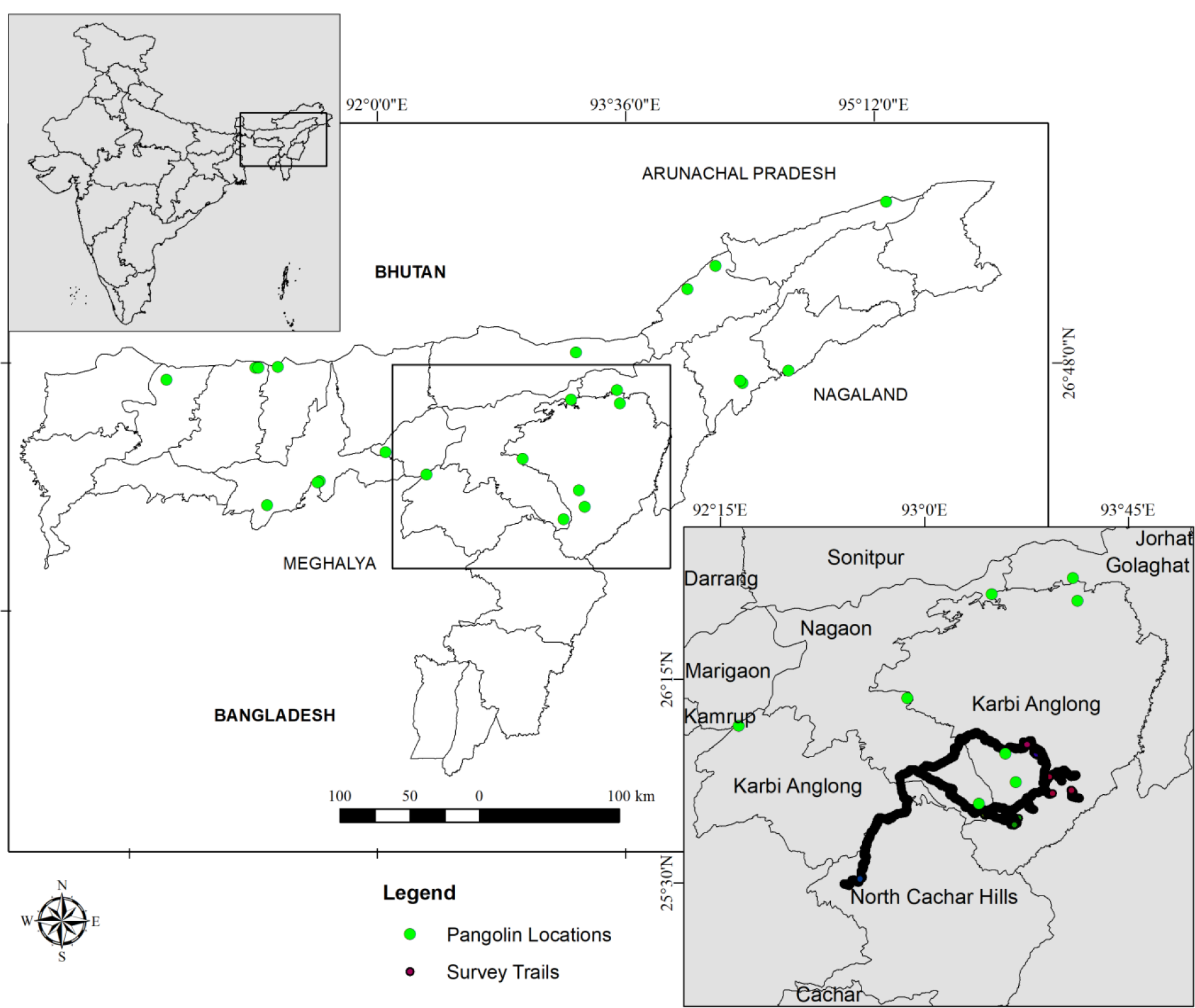
Map of the study area showing survey tracks/trails (imported from GPS device) and pangolin GPS locations collected through direct and indirect sources in Assam, India

**Fig.2.**
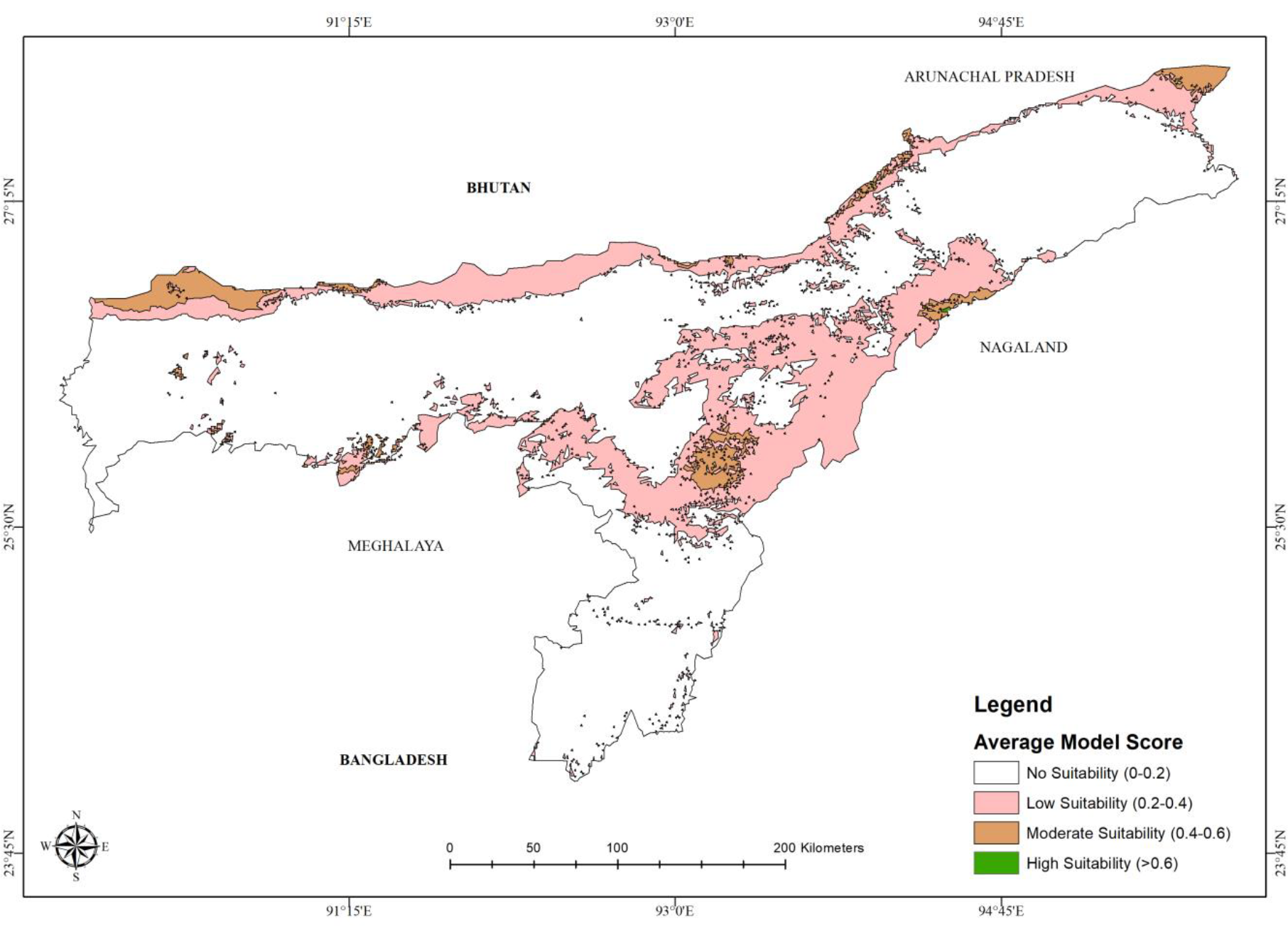
Suitable habitat thresholds of Chinese pangolin in Assam, India predicted by the two models (mean of the models)

**Fig. 3.**
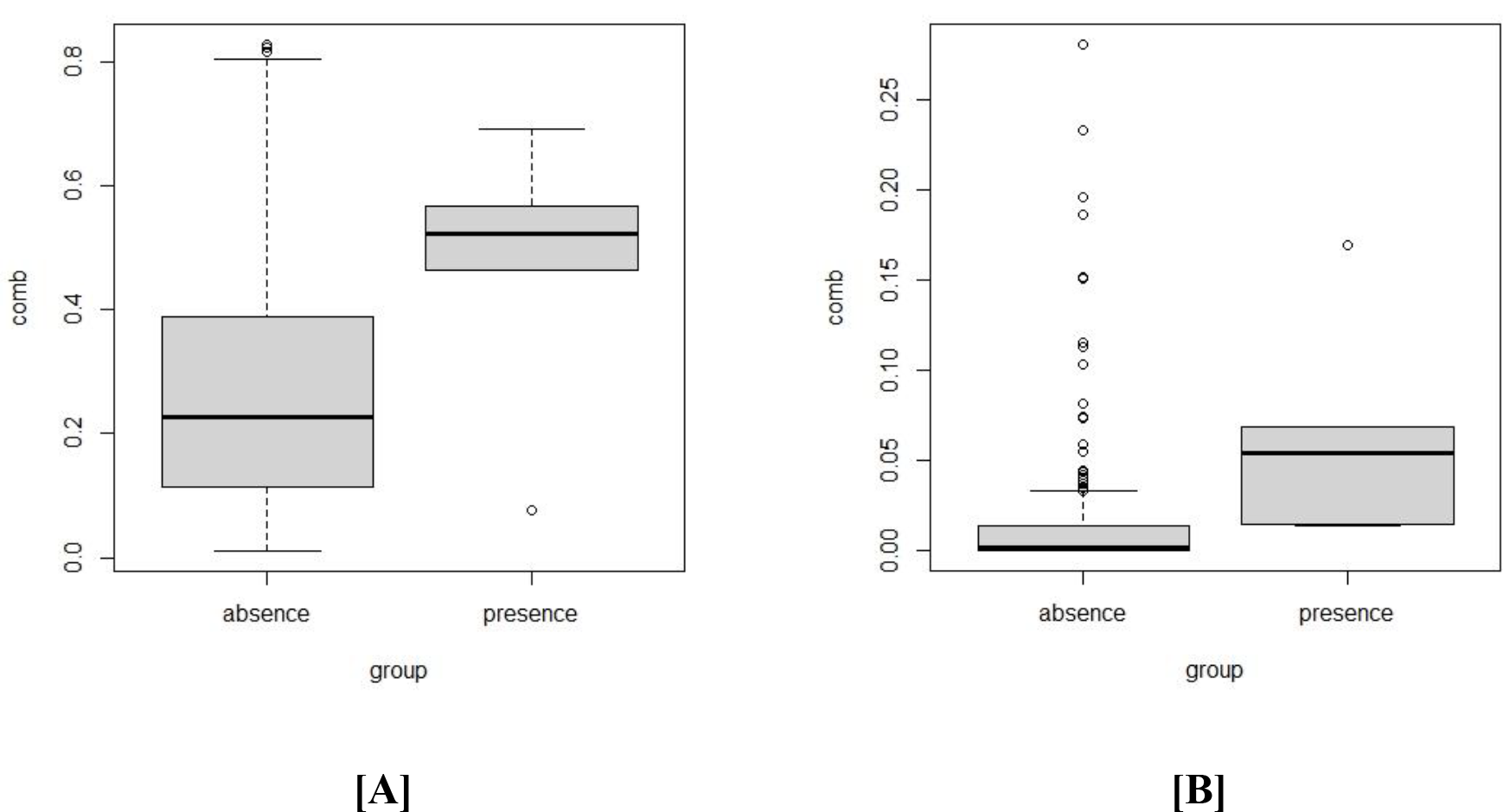
Boxplot showing [A] Maxent and [B] Random Forest presence absence data points

**Fig. 4.**
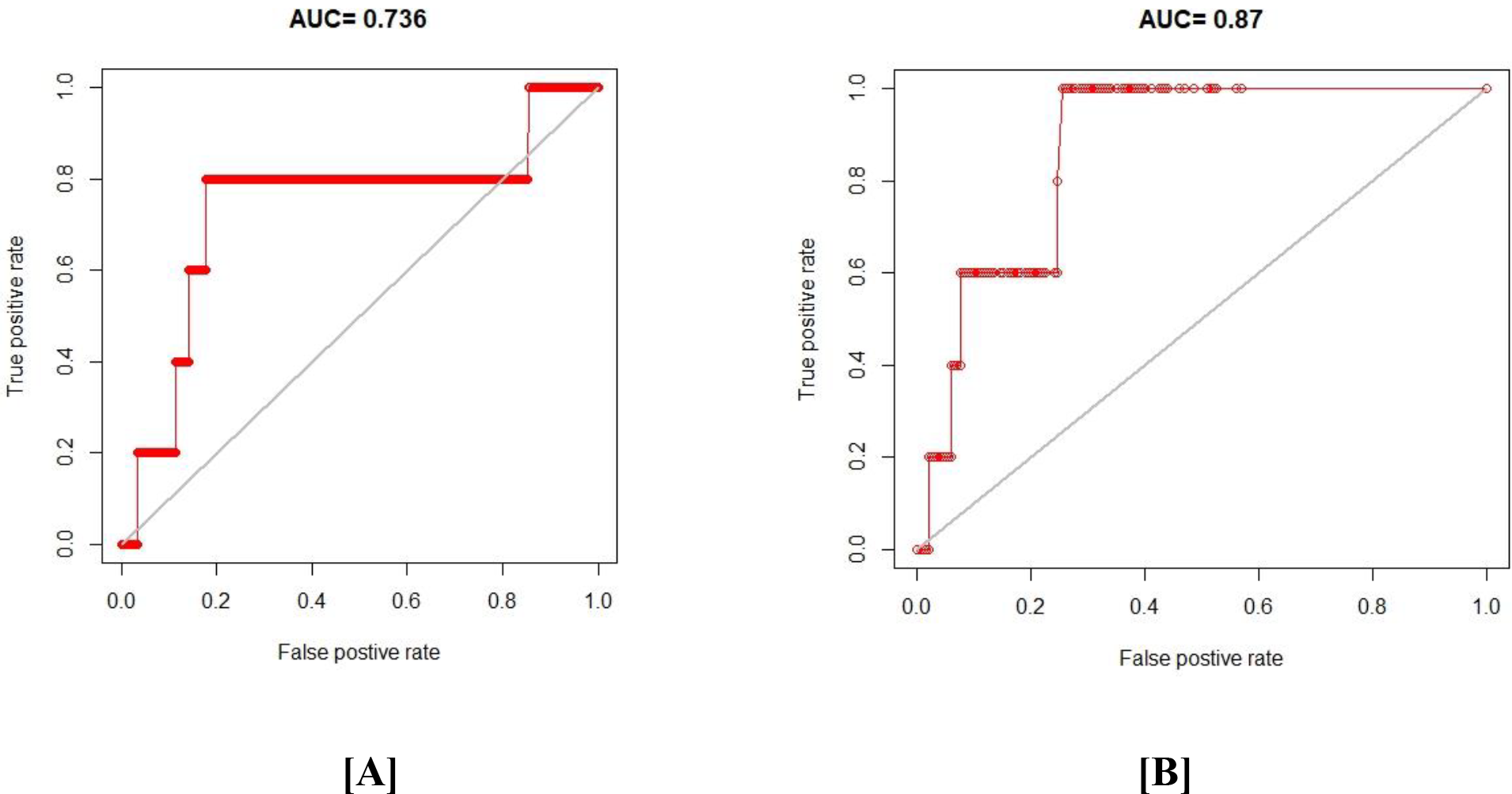
ROC comparison between [A] Maxent and [B] Random Forest model

Variations in the predicted distribution of Chinese Pangolin in Assam have been seen in the individual model outputs (Fig. 5). The Maxent model projected that more would be within a different level of suitability, while the Random Forest forecasted very low territory. On the other hand in Karbi Anglong district of Assam, very low percentage area was found under the high suitability area (). Highest percentage area was found under low suitability threshold () followed by moderate suitability () (Fig. 7).

**Fig.5.**
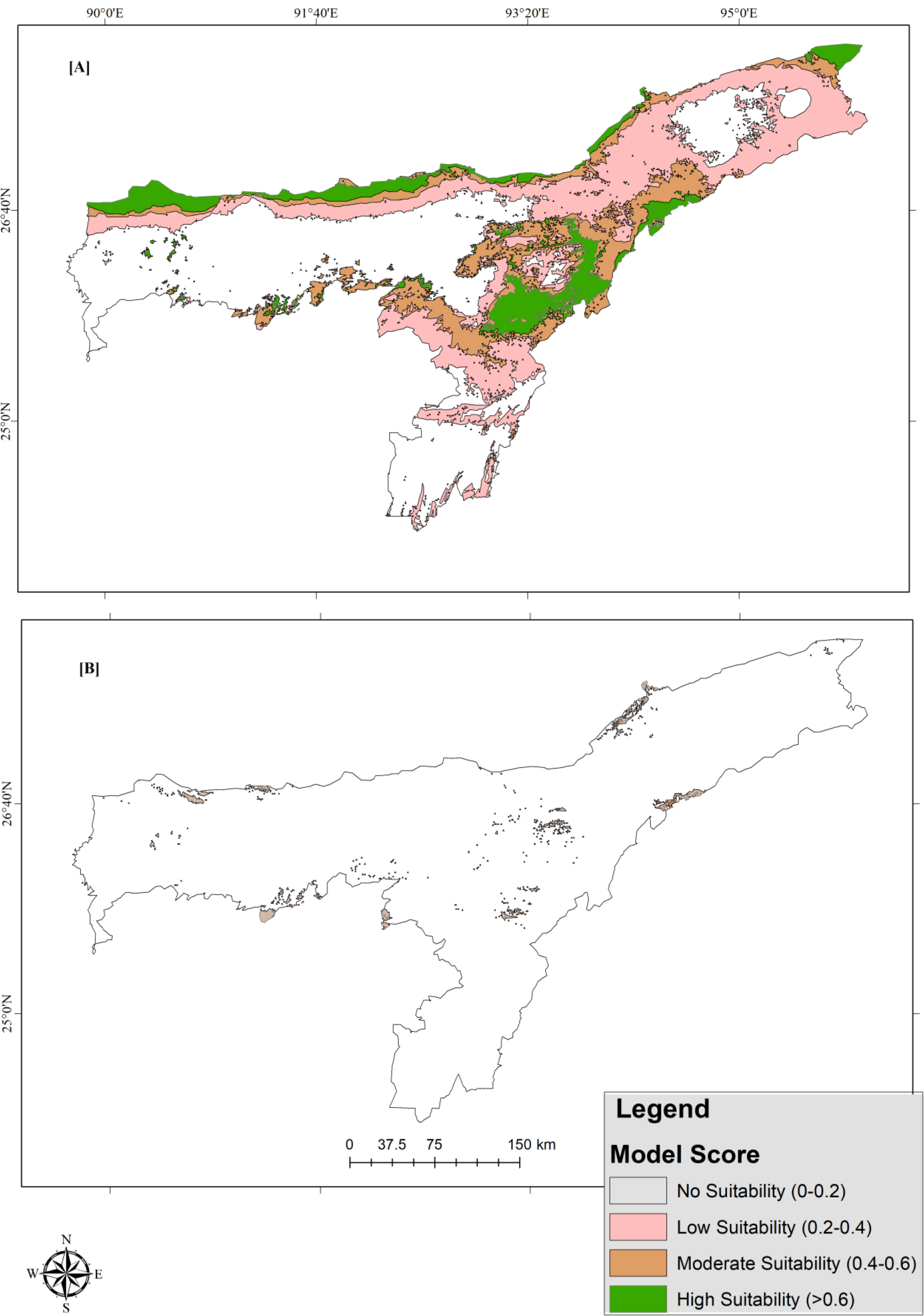
Suitable habitat map of Chinese pangolin in Assam, India predicted by [A] Maxent model and [B] Random Forest model.

**Fig. 6.**
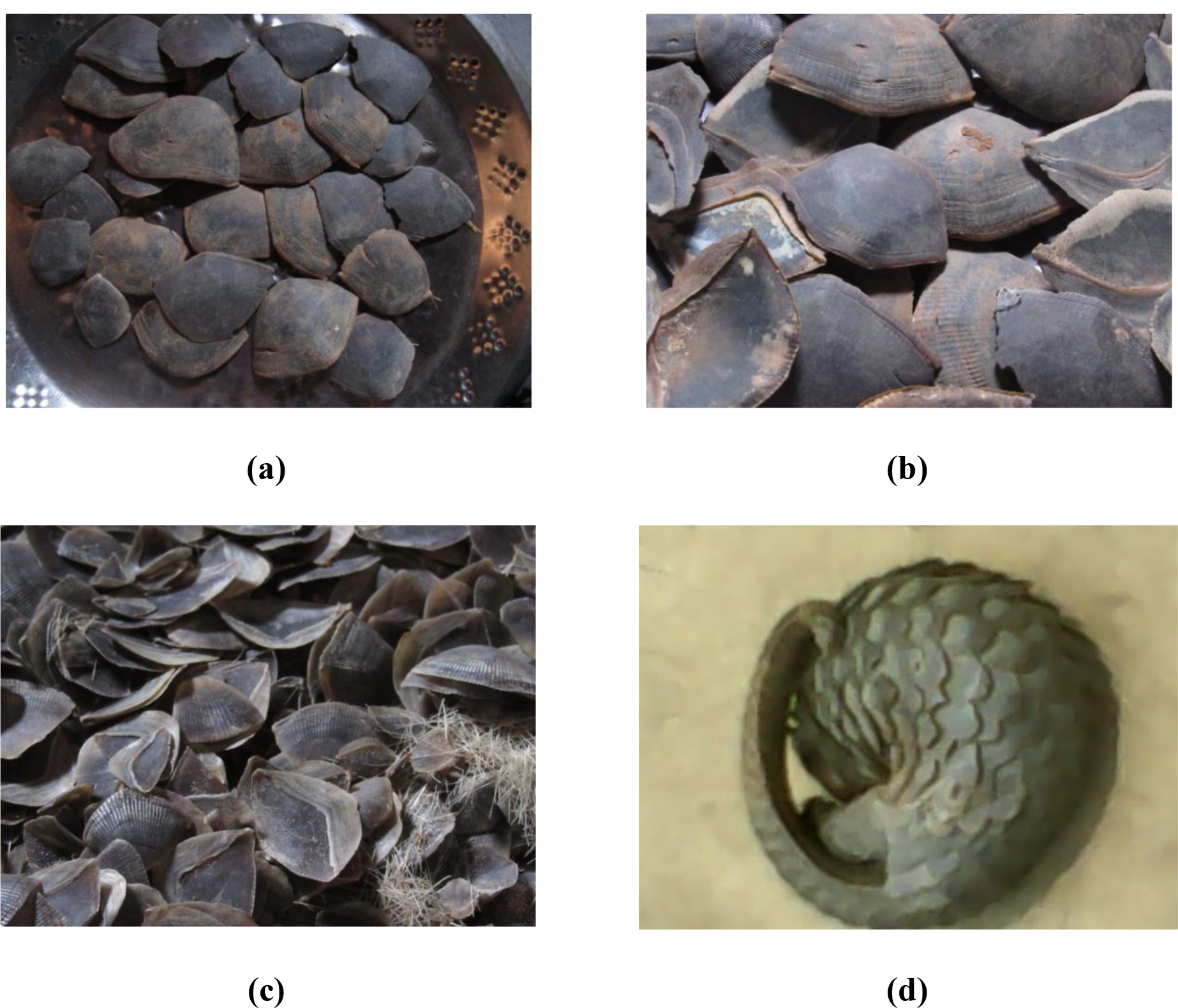
Photographs collected during the study period: (a), (b), (c) scales of pangolin kept in households;(d) a dead pangolin

**Fig.7.**
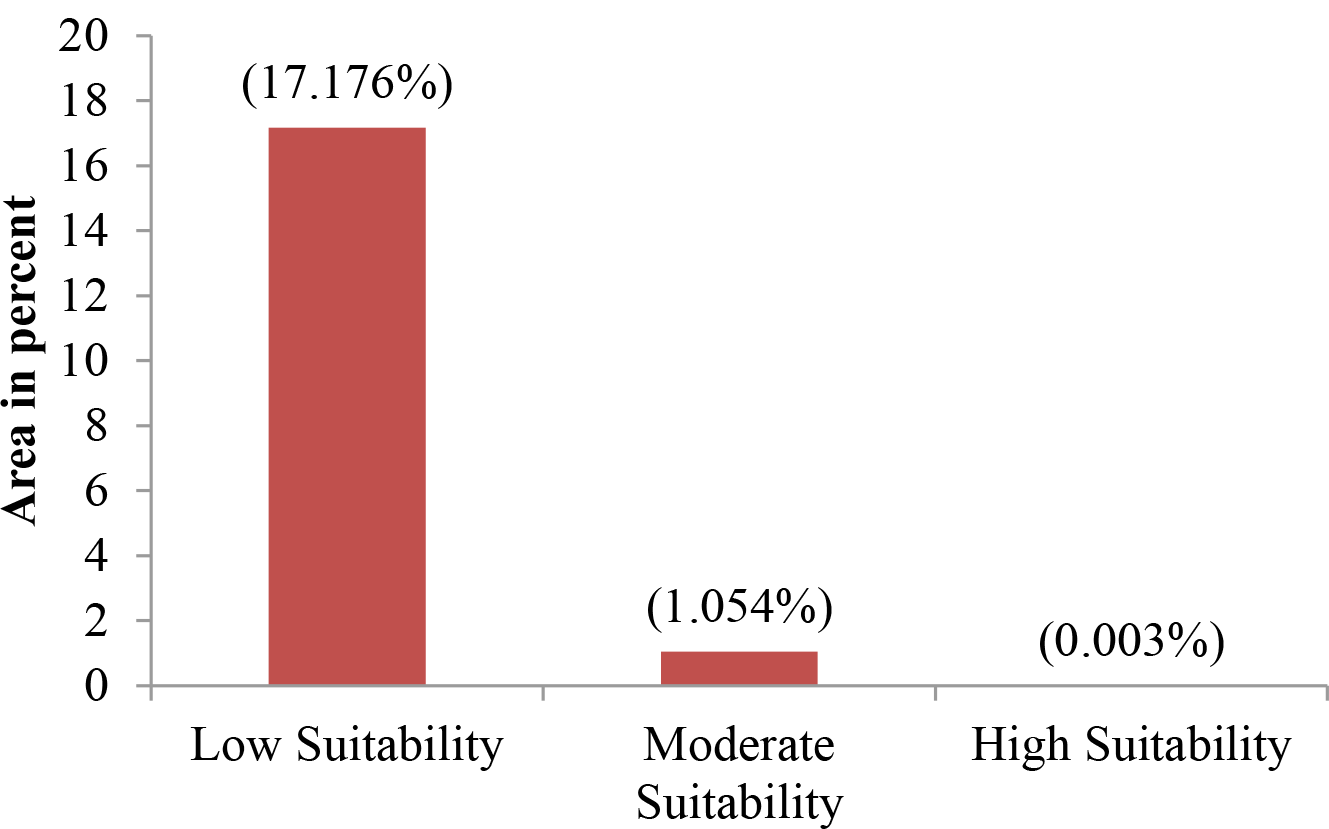
Percentage area under suitability threshold derived from average model output map within Karbi Anglong District (No suitability area is excluded)

The past distribution record of Chinese pangolin revealed from the household survey revealed that pangolins were present in their vicinity before more than 10 years. The species has been least seen by most of the respondents in the last 10 years (43.7%). Most of the respondent of age class 18-24 years never saw a pangolin (82.5%) in the area which depicted the population decline of the spcies. The species is mostly hunted in accidental encouter (23.5%). 49% of the respondents has ate the meat of the spcies through direct or indirect sources. Moreover, low percentage of respondents were agreed to the medicinal use of the species (21.2%) and claimed that they did not know about the medicinal use of the species (78.8%). Scales were the most used body parts for traditional medicine (58.8%) especially for exceesive saliva secretion in children. Most of the respondents who knows about chinese pangolin (66.2%) also commented on the burrows of the species.

The tree canopy cover loss area was found highest in the year 2014 (41.97 ha), 2009 (1545.98 ha) and 2016 (9970.218 ha) for high, moderate and low suitable habitat respectively (Figure 8). The increasing trend of tree canopy cover was found in high suitable habitat (Sen’s slope = 0.49, z=2.309, p<0.05), low (Sen’s slope = 180.13, z= 1.82, p>0.05) and moderate suitable habitat (Sen’s slope=9.59, z=0.84, p>0.05).

**Fig. 8.**
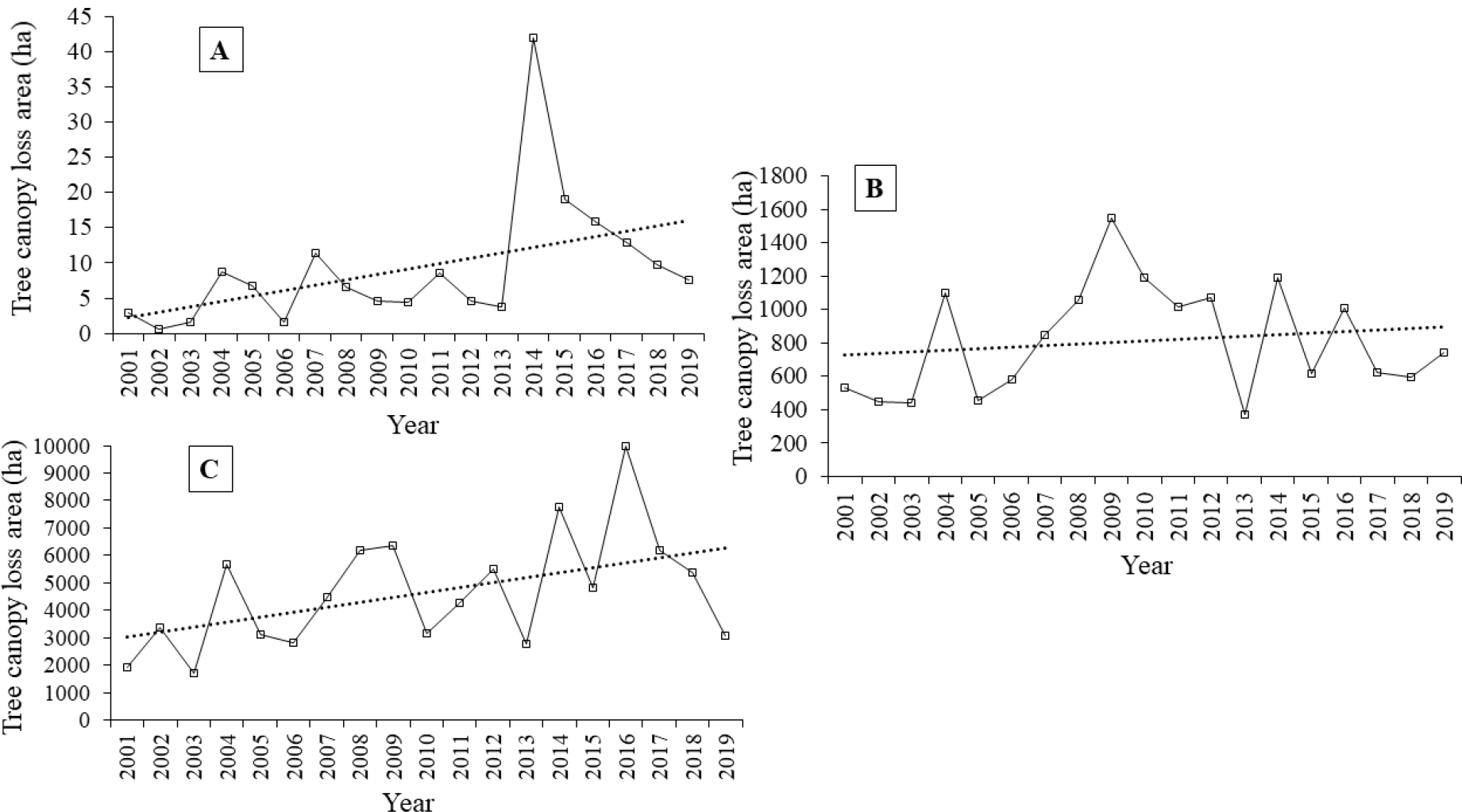
Annual tree cover change (with linear trend); A. High suitable habitat, B. Moderate suitable habitat and C. Low suitable habitat

## DISCUSSION

In view of the global population decline of Chinese pangolin, the high priority of studying the habitat interactions of this species is urgently required. The little known natural history of the species makes it more difficult, particularly in terms of the collection and quantification of ecological data. Much of the Chinese pangolin’s ecological data derives from studies in Taiwan and southern China (e.g., Lu, 2005; Lin, 2011). In Indian context, Chinese pangolin is least studied species owing to its low delectability (direct sightings). Moreover, due to overexploitation of Asian species including Chinese pangolin, very low population densities are expected resulting difficulties in surveying the species (Wilcox et al., 2019). This is evident in populations of Asian species in particular, where decades of targeted and sustained over-exploitation have resulted in very low population densities (e.g., Duckworth et al., 1999).

The results of the present study provide detailed information of spatial distribution of Chinese pangolin determined by various categories of habitat suitability in the state of Assam. The importance of the environmental attributes has been well depicted by the two machine learning methods applied to predict the suitable habitat of the species. In addition, the efficacy of the models has been illustrated through various parameters such as AUC, Kappa coefficient etc. Altitude and seasonality of temperature are two major factors influencing the distribution of Chinese pangolin in Assam. The Random Forest model has shown higher AUC value than that of Maxent model supporting the assumption that the species has a rather limited distribution range as the AUC values appear to be lower for species with broad distribution (Ardestani et al. 2015; ZareChahouki and PiriSahragard 2016). Further, the relationship between the predicted map and the actual distribution map is well represented by the Kappa coefficient and provides improved results for Maxent (0.464) than that of Random Forest (0.013). Kappa coefficient is the sum of true positive rate (TPR) and true negative rate (TNR) determined from the confusion matrix of the model. However, the performance of the model is strongly correlated with the species being modelled and it is impossible to find a superior model based on its performance (PiriSahragard and ZareChahouki 2016).

High potential suitable habitat in the foot hill region of the state of Assam has also been supported by the encounter data of Chinese pangolin in the region. A similar result of high frequency of encounters between people and Chinese pangolins has been reported from Nepal in forested areas in the hills and mountain regions (Gurung, 1996; Reddy et al., 2018; Katuwal et al., 2017). In addition, the human population growth and expansion of agricultural land is posing high threats to the foothill forests particularly in the northern boundary of the state of Assam. However, the model output needs extensive field verification to provide a conclusive idea about the habitat of the species. There may be many other factors that affect the abundance of the species other than habitat loss and hunting such as availability of the prey, seasonality and predation. It is evident from our result that the tree cover in the high suitability area has been decreasing during 2001 to 2019. This may contribute in drastic change in the ecosystem leading to change in the microclimate. Although, microclimate of the burrows may not be altered in the process, the food availability may be significantly influenced (Bao et al., 2013; Mahmood et al., 2013).

The traditional ecological knowledge of the species is found to be very poor in the study area and is greatly biased among age class. That also depicts low detestability of the species. But the major concern in the study area is accidental encounter of the species in which most of the individuals of the species are killed. Although, the respondents were reluctant to share hunting information, 49% of them have eaten the meat of the species from direct or indirect sources. The Karbi tribe believes that encountering Chinese pangolins is as bad omen as it would bring famine to society. They have also argued that the pangolins are the descendants of a new born baby who died in starvation in the countryside, and that is why the pangolin has no teeth. However, very little information about the illegal trade and their use in traditional medicine has been generated in the present study.

## Acknowledgements

The authors express their sincere thanks to Principal Chief Conservator of Forest, KarbiAnglong Autonomous Council (KAAC), Assam for giving necessary permission to carry out the research and Conservation Leadership Programme (CLP) and Rufford Small Grant Foundation for generous financial support. Authors are also thankful to Biranjoy Basumatary and Rustom Basumatary for their assistance in field.

## Notes

### Competing Interest Statement

The authors have declared no competing interest.

